# Bioengineering a light-responsive encapsulin nanoreactor: a potential tool for photodynamic therapy

**DOI:** 10.1101/2020.06.06.138305

**Authors:** Dennis Diaz, Xavier Vidal, Anwar Sunna, Andrew Care

**Author notes:** Corresponding author: Andrew Care, Department of Molecular Sciences, Macquarie University, Sydney, NSW 2109, Australia.

## Abstract

Encapsulins, a prokaryotic class of self-assembling protein nanocompartments, are being re-engineered to serve as ‘nanoreactors’ for the augmentation or creation of key biochemical reactions. However, approaches that allow encapsulin nanoreactors to be functionally activated with spatial and temporal precision is lacking. We report the construction of a light-responsive encapsulin nanoreactor for “on-demand” production of reactive oxygen species (ROS). Herein, encapsulins were loaded with the fluorescent flavoprotein mini-Singlet Oxygen Generator (miniSOG), a biological photosensitizer that is activated by blue-light to generate ROS, primarily singlet oxygen (^1^O_2_). We established that the nanocompartments stably encased miniSOG, and in response to blue-light were able to mediate the photoconversion of molecular oxygen into ROS. Using an *in vitro* model of lung cancer, ROS generated by the nanoreactor was shown to trigger photosensitized oxidation reactions that exerted a toxic effect on tumour cells, suggesting utility in photodynamic therapy. This encapsulin nanoreactor thus represents a platform for the light-controlled initiation and/or modulation of ROS-driven processes in biomedicine and biotechnology.

## Introduction

In the era of synthetic biology, there is increasing interest in re-engineering sub-cellular compartments (e.g. organelles) into designer ‘nanoreactors’ that can augment/enhance existing biological reactions, or enable the creation of entirely new synthetic reactions.^1-3^ This includes protein compartments that encase enzymes within a selectively permeable protein shell that self-assembles from multiple protein subunits. ^4, 5^ This design isolates and promotes specialized reactions by placing catalytic proteins, their substrates and cofactors within close proximity of each other; regulating the influx and efflux of molecules; preventing the escape of volatile or toxic reaction intermediates; and creating distinct microenvironments that can protect, stabilise and improve the function of cargo proteins. ^6^

Encapsulins are a newly established class of prokaryotic protein nanocompartments, which self-assemble from identical protein subunits into semi-permeable icosahedral protein shells with *T* = 1 (60 subunits, 20-24 nm), *T* = 3 (180 subunits, 30-32 nm) or *T* = 4 (240 subunits, 43 nm) symmetries. ^3, 7-9^ In nature, they encase functional cargo proteins that help their microbial hosts maintain iron homeostasis, cope with oxidative and nitrosative stress, and/or safely derive energy from ammonium. ^10-12^ Moreover, encapsulins recognise and selectively encapsulate native cargo proteins that display a short encapsulation signal peptide (ESig), a distinct mechanism that can be adapted to instead package foreign cargo, reprogramming the nanocompartments’ functionality. ^7, 12, 13^

The structural modularity, functional programmability and widespread prevalence of encapsulins in prokaryotes, makes them an ideal toolbox for nanoreactor construction. By packaging non-native enzymes, encapsulins have been recently reprogrammed to stabilize the synthesis of precursors used in the production of pharmaceutical opioids; and to safely compartmentalize the orthogonal synthesis of toxic melanin thereby mimicking melanosome organelles found in mammals.^1, 2^ However, the capacity to selectively control the activity of such encapsulin nanoreactors is lacking. Herein, we explore the construction of an encapsulin nanoreactor that generates reactive oxygen species (ROS) in response to external light stimulation, permitting precise temporal and spatial control over its activity.

As a first step towards designing a light-responsive encapsulin nanoreactor, we elected to load nanocompartments with mini-Singlet Oxygen Generator (miniSOG). miniSOG is a flavin mononucleotide (FMN)-based fluorescent protein derived from the light-oxygen-voltage-sensing domain of *Arabidopsis thaliana* phototropin-2. Via its FMN chromophore, miniSOG can be activated by blue-light to photoconvert molecular oxygen (^3^O_2_) into ROS, primarily singlet oxygen (^1^O_2_). ^14^ Consequently, in a process called ‘photosensitization’, ^1^O_2_ generated by light-induced miniSOG has been exploited in various applications. These include the photooxidation of contrast agents in bioimaging, photoredox activation of prodrugs, photomodulation of ROS-activated signalling pathways, chromophore-assisted light inactivation of proteins in optogenetics, and photoinduced destruction of cancer cells in photodynamic therapy (PDT). ^15-22^

In this communication, we present the confinement of ESig-tagged miniSOG inside *Thermotoga maritima* encapsulin, reprogramming it to serve as a light-responsive nanoreactor for “on demand” ROS generation. We show the capacity of this first-of-its-kind encapsulin nanoreactor to house miniSOG and its blue-light induced photoconversion of ^3^O_2_ to _^1^_O_2_. We further characterize the effect encapsulation has on miniSOG’s ^1^O_2_-generating function, and the impact of prolonged ROS generation on the nanocompartments’ structure and stability. As a proof-of-concept, the ROS produced by the light-activated nanoreactor is demonstrated to trigger photosensitized oxidation reactions that exert a phototoxic effect on lung cancer cells, indicating potential utility in PDT.

## Materials and methods

### Materials

All chemicals and reagents used in this study were purchased from Sigma-Aldrich unless stated otherwise.

### Molecular biology and cloning

All inserts were codon optimized for expression in *Escherichia coli* and custom synthesized as gBlock Gene Fragments (Integrated DNA Technologies). A surface-exposed loop region between residues 138 and 139 of the encapsulin from *Thermotoga maritima (Tm)* (Uniprot: Q9WZP3) was modified with a hexahistidine tag (GGGGGGHHHHHHGGGGGG) (his-tag) as previously described. ^23^ For the encapsulation of miniSOG inside his-tagged encapsulin (Enc), the protein was C-terminally tagged with a minimized *Tm* encapsulation signal (ESig, GGSENTGGDLGIRKL), ^24^ resulting in ESig-tagged miniSOG (mSOG).^25^ To generate expression vectors, *Enc* was ligated into pETDuet-1 (Merck, USA) via NcoI/BamHI restrictions sites, while *mSOG* was inserted into pACYC-Duet-1 (Merck, USA) via NdeI/BglII restriction sites. *Escherichia coli α*-Select (Bioline, UK) was used as a host for plasmid propagation. Gene insertion was verified by PCR using the primer pairs pETUpstream/DuetDOWN or DuetUP2/T7 Terminator (Merck) with Enc or mSOG-containing plasmids as template DNA. All plasmids constructed for this study are summarized in Supplementary Table S1. *Escherichia coli* BL21 (DE3) cells (New England Biolabs, USA) were used for recombinant protein expression. For the co-expression of Enc and its intended mSOG cargo protein, cells were co-transformed with the appropriate expression plasmids, and the resulting transformants were selected on Luria–Bertani (LB) agar supplemented with carbenicillin (100 μg/ml) and chloramphenicol (50 μg/ml) (see Supplementary Table S2).

### Protein expression and purification

Protein expression (or co-expression) experiments were carried out in LB medium supplemented with carbenicillin (100 μg/ml), chloramphenicol (50 μg/ml), or both. Briefly, 500 ml LB was inoculated 1:100 with an overnight culture of cells harbouring the expression plasmid(s) of interest. Then, the culture was grown aerobically at 37 °C to an optical density at 600 nm (OD_600_) of 0.5-0.8 and induced by the addition of isopropyl-ß-D-thiogalactopyranoside (IPTG). The optimized conditions for the recombinant expression of all proteins in this study are outlined in Table S2. Finally, cells were harvested by centrifugation (10,000 × *g* 15 min, 4 °C) and stored at −30 °C. For the purification of Enc and mSOG-loaded Enc (Enc-mSOG), the pellet from a 500 mL culture was resuspended in 50 mL lysis buffer (20 mM NaH_2_PO_4_, 300 mM NaCl, 40 mM imidazole, 1 U/mL Benzonase nuclease, pH 7.4). Cells were lysed with a French pressure cell press and subsequently centrifuged at 10,000 × *g* for 15 min. The lysate was then subjected to nickel-immobilized metal affinity chromatography (IMAC) using a HisPrep™ Fast Flow 16/10 column (GE Healthcare, USA) equilibrated with equilibration buffer (20 mM NaH_2_PO_4_, 300 mM NaCl, 40 mM imidazole, pH 7.4). The imidazole concentration within the equilibration buffer was increased to 260 mM or 400 mM to elute Enc-mSOG and Enc, respectively. Next, eluted protein fractions were concentrated using Amicon® Ultra-15 centrifugal filter units (Merck, USA) with a 100 KDa cut-off, followed by dilution in 7 mL of 50 mM HEPES buffer pH 7.4 (Chem-Supply Pty, Australia). A second purification step by size exclusion chromatography (SEC) was subsequently performed using a HiPrep™ 26/60 Sephacryl® S-500 HR column (GE Healthcare, USA) and 100 mM HEPES Buffer. All purifications were carried out on an Äkta™ start or Äkta™ pure chromatography system (GE Healthcare, USA). For the purification of free mSOG the unbound fraction obtained during the IMAC purification of Enc-mSOG was used. Herein, a saturated solution of ammonium sulphate was added to a final concentration of 30% (v/v), incubated on ice for 30 min and spun down at 10,000 × *g* for 15 min. Next, ammonium sulphate was added to the supernatant to a final concentration of 50% (v/v), inducing protein precipitation. The precipitated protein was resuspended in 100 mM HEPES buffer (pH 7.4) and subjected to SEC using a HiPrep™ 16/60 Sephacryl® S-400 column (GE Healthcare, USA) using 100 mM HEPES Buffer. The fractions containing free mSOG were pooled and concentrated using Amicon Ultra-15 centrifugal filter units with a 10 KDa cut-off. The final protein concentrations were determined by measuring absorbance at 280 nm. Examples of purification chromatograms are provided in Supplementary Figure 1.

### Polyacrylamide gel electrophoresis (PAGE)

The Bio-Rad mini-protean system (Bio-Rad laboratories, USA) was used for all SDS-PAGE and Native-PAGE analysis. For SDS-PAGE, samples were diluted in 2X Laemmli sample buffer with 50 mM 1,4-dithiothreitol, heated at 95 °C for 5 min, loaded into pre-cast Bio-Rad Mini-PROTEAN® TGX™ gels (4-15 %) and run at 200 V for 30 min. For Native-PAGE, samples were diluted in 4X native sample buffer (200 mM Tris-HCl pH 6.8, 40% glycerol, and 0.08% bromophenol blue), loaded into pre-cast Bio-Rad Mini-PROTEAN® TGX™ gels (4-20%) and run at 200 V for a minimum of 2 h. In-gel fluorescence of proteins was observed with a gel documentation imager (Bio-Rad laboratories, USA). All gels were stained following the Coomassie G-250 safe stain protocol ^26^. The densitometric intensity of protein bands in SDS-PAGE gels were quantified using ImageJ software (National Institutes of Health, USA) ^27^.

### Transmission electron microscopy (TEM)

10 µL of Enc or Enc-mSOG (∼100 µg/ml) was adsorbed onto formvar-carbon coated copper grids for 2 min and negatively stained with uranyl acetate replacement stain (UAR-EMS) for 1 h. Grids were then washed with ultrapure water and allowed to dry for 15 min. Finally, the grids were observed under a CM10 TEM (Philips, Netherlands) operated at 100 kV accelerating voltage.

### Dynamic light scattering (DLS)

DLS data was collected on a Nano ZS90 Zetasizer (Malvern, UK). Measurements were performed at room temperature using standard cuvettes containing 1 mL of Enc or Enc-mSOG were diluted in 100 mM HEPES (pH 7.4) to a final concentration 0.2-0.4 mg/mL. The signal was averaged over 13 readings, each lasting 30 s.

### Absorbance and fluorescence spectrometry

The fluorescence excitation and emission spectra of free and encapsulated mSOG were obtained on a Cary Eclipse Fluorescence Spectrophotometer (Agilent Technologies, USA) or Fluorolog® (Horiba, Japan) using quartz cuvettes. Absorbance for protein concentration measurements was acquired at 280 nm on a SPECTROstar*®* Nano Plate Reader (BMG Labtech, Germany) using UV-transparent 96 well plates.

### Singlet oxygen detection

Singlet oxygen generation from free mSOG, Enc-mSOG, and unloaded Enc was detected in solution with the fluorescent probe Singlet Oxygen Sensor Green (SOSG) according to the manufacturer’s protocol (Invitrogen, USA). The reaction mixture contained 500 nM of free mSOG or encapsulated mSOG (Enc-mSOG) or 4.8 of µM unloaded Enc (in 100 mM HEPES buffer, pH 7.5), 1 µM SOSG, and 50% deuterium oxide (D_2_O). The concentration of unloaded Enc is equivalent to the concentration of the nanocompartment shell present in Enc-mSOG reactions. Reaction mixtures were irradiated with a Chameleon-Ultra II laser (Coherent) passing through a harmonic converter to a set wavelength of 450 nm with an average power density of 55 mW/cm^2^ for 10 min. For further characterization, other irradiation times were evaluated (0, 10, 15 and 20 min). Fluorescence signals from the oxidized SOSG (excitation/emission = 485/520 nm) were measured on a PHERAstart FS (BMG Labtech, Germany) microplate reader.

### In vitro cytotoxicity and phototoxicity

For *in vitro* cytotoxicity and phototoxicity studies, 5.0 × 10^3^ A549 cells per well were seeded into 96-well microplates and cultured at 37 °C for 24 h. First, the effect of laser irradiation on cell viability was investigated by exposing cells to a 450 nm blue laser at a power density of 55 mW/cm^2^ for different time periods (0, 5, 10 and 15 min). To evaluate the cytotoxicity of all protein constructions in the absence of light activation, cells were incubated with 4.8 µM unloaded Enc and 500 nM mSOG or Enc-mSOG at 37 °C in the dark for 2, 4, 8 and 12 h. After being subjected to each of these treatments, cells were washed once with PBS to remove non-internalized protein, and fresh growth medium was added. Cells were cultivated for a further 48 h, and cell viability was then determined using the 3-(4,5-dimethylthiazol-2-yl)-2,5-diphenyltetrazolium bromide (MTT) cell viability assay (Invitrogen, USA) according to the manufacturer’s protocol ^28^. In phototoxicity studies, the same protocol described for cytotoxicity was performed with minor changes. Briefly, after A549 cells were treated with free mSOG, Enc-mSOG, and unloaded Enc for different times (2, 4, 8 and 12 h) the medium was replaced with PBS and cells were irradiated with a 450 nm blue laser at 55 mW/cm^2^ for 10 min. Next, fresh medium was added to the cells, followed by cultivation for another 48 h in the dark. Cell viability was subsequently measured by MTT assay. For each experiment at least three technical replicates were performed.

## Results and discussion

### Reprogramming an encapsulin nanocompartment into a light-activatable nanoreactor

In nature, *Thermotoga maritima* (*Tm*) encapsulin packages a ferritin-like protein that facilitates the sequestration, oxidization and biomineralization of iron, indicating a physiological role in iron homeostasis and oxidative stress responses. We aimed to bioengineer a light-activatable nanoreactor for the production of ROS by reprogramming the *Tm* encapsulin to encase the photosensitizing protein miniSOG. To selectively target miniSOG’s encapsulation inside *Tm* encapsulin, its C-terminus was tagged with a functional minimized 15-amino acid *Tm* ESig. ^25^ For the heterologous production of mSOG-loaded encapsulin (Enc-mSOG) in *E. coli*, the ESig-tagged miniSOG (mSOG) cargo was co-expressed with a His-tagged encapsulin (Enc) (Figure 1a). Following their purification by IMAC and SEC, both Enc-mSOG and unloaded Enc underwent biophysical characterization (Figure 1b-e). SDS-PAGE confirmed the co-purification of Enc (Enc_subunit_; 31.9 kDa) and mSOG cargo (mSOG-ESig; ∼15.9 kDa) (Figure 1b, left panel).Densitometric SDS-PAGE gel analysis determined the ratio of Enc_Subunit_ to mSOG cargo and estimated a cargo loading capacity (LC%) of 7-9% for Enc-mSOG, which represents ∼7 ± 2 mSOG molecules packaged per nanocompartment. Under native-PAGE conditions, Enc-mSOG presented high molecular weight bands similar to unloaded Enc, consistent with *T*=1 Enc assemblies (Figure 1b, right panel). The blue-light excitation of mSOG proteins inside Enc-mSOG was detected via fluorescence imaging of the native-PAGE, with no fluorescence observed from empty Enc (Figure 1b, right panel). In order to confirm the correct self-assembly, morphology and size of mSOG-loaded nanocompartments, TEM observations and DLS measurements were performed. TEM images of negatively stained samples showed the accurate formation of Enc-mSOG and empty Enc into spherical nanocompartments (Figure 1c, upper panel). DLS measurement of empty Enc revealed a mean hydrodynamic diameter of 29.0 ± 2.5 nm (Figure 1c, lower panel), which was expected to be ∼24 nm based on the crystal structure of *Tm* encapsulin (*T*=1; Protein database ID: 3DKT). ^25^ This observed enlargement is likely due to the insertion and display of His-tags on the Enc’s external surface. This aligns with research by Moon et al, in which *Tm* encapsulin enlarged to 29.1 nm after introducing functional peptides into the same 138-139 loop region ^23^. Nevertheless, DLS determined that unloaded Enc and Enc-mSOG were both intact and monodisperse with mean diameters of ∼30 nm, thus mSOG packaging did not significantly alter Enc’s morphology or structure.

**Fig. 1.**
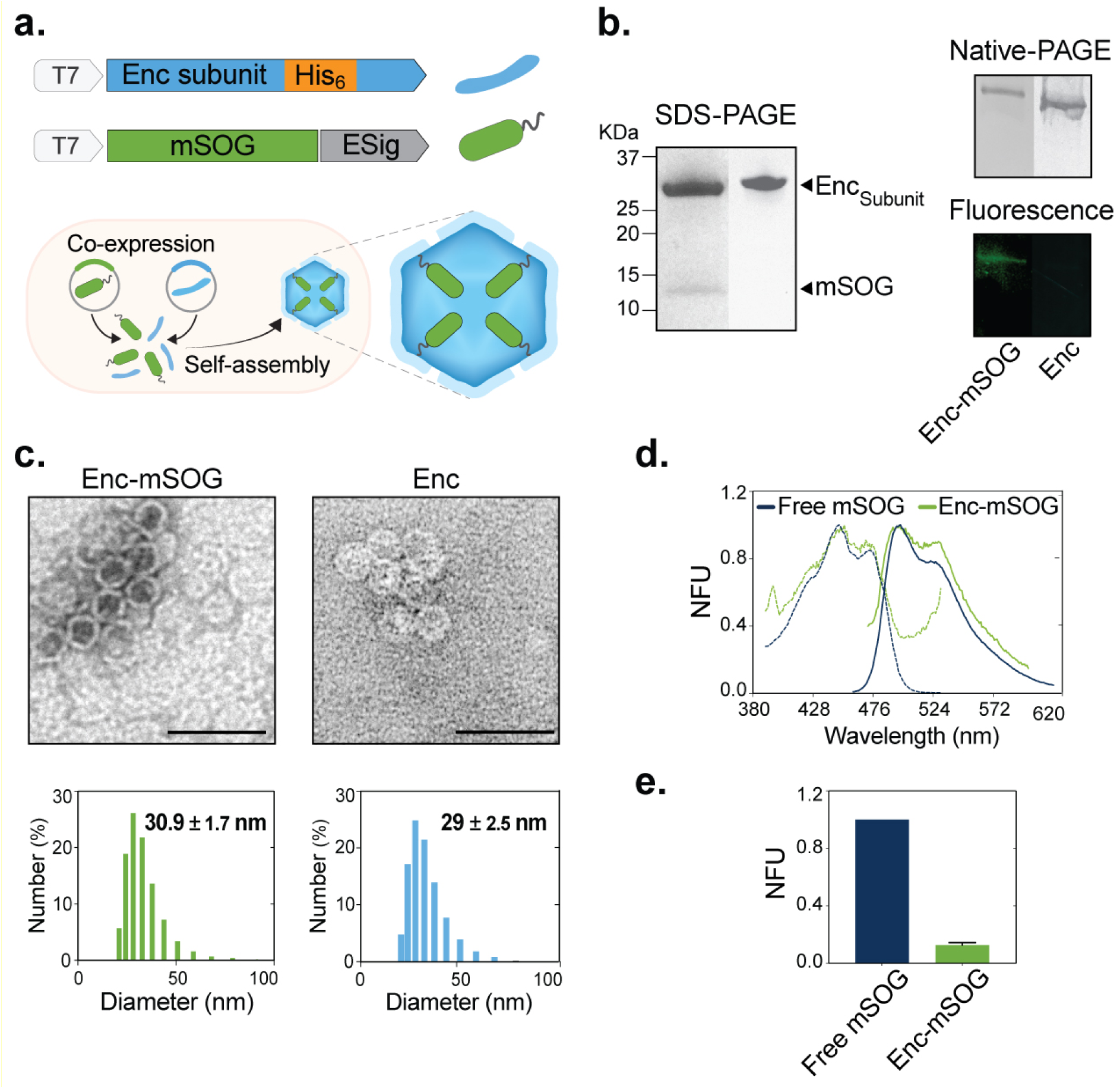
Design, production and biophysical characterization of mSOG-loaded encapsulin (Enc-mSOG). **(a) (Upper panel)** Genetic constructions encoding the Enc subunit from *Thermotoga maritima* (*Tm*) (blue) displaying a His-tag (orange) within a surface-exposed loop region, and mSOG cargo protein (green) C-terminally tagged with an encapsulation signal (ESig, grey). **(Lower panel)** Heterologous co-expression of encapsulin subunits and ESig-tagged mSOG in *E. coli* leads to the *in vivo* self-assembly of cargo-loaded *Tm* Enc *T*=1 nanocompartments. **(b)** PAGE analysis of Enc-mSOG purified by sequential IMAC and SEC. **(Left panel)** SDS-PAGE showing the co-purification of the Enc subunit (31.9 KDa) and ESig-tagged mSOG cargo protein (14.4 KDa). **(Right panel)** Native-PAGE verifying the self-assembly of Enc-mSOG into cargo-loaded nanocompartments, and in-gel fluorescence of the Native-PAGE confirming encapsulation of fluorescent mSOG cargo. **(c) (Upper panel)** TEM images of Enc-mSOG and unloaded Enc show their self-assembly into spherical nanocompartments (scale bars = 50 nm). **(Lower panel)** While their respective size distributions measured by DLS indicate average diameters of ∼30nm. **(d)** Effect of encapsulation on the fluorescence excitation/emission spectra of mSOG. Fluorescence excitation (dashed line) and emission (solid line) spectra of free mSOG and Enc-mSOG given in Normalized Fluorescence Units (NFU). **(e)** Fluorescence emission intensity (ex/em = 485/520 nm) of free mSOG and Enc-mSOG. Each sample contained 500 nM mSOG equivalent. Error bars represent the mean ± standard deviation, n=3 from three independent experiments.

Next, spectrophotometric analysis was used to gain insight into the effect encapsulation had on the fluorescent properties of mSOG. According to Figure 1d, free mSOG and Enc-mSOG have almost identical fluorescence excitation maxima at 450 nm (with shoulders at 470 nm) and emission maxima at 495 nm (with shoulders at 525 nm). These spectra are consistent with the reported fluorescence spectra of unmodified miniSOG. ^24^ However, we observed that mSOG’s fluorescence spectra became nosier upon encapsulation and coincided with an 87% reduction in its fluorescence intensity (Figure 1e). This marked loss of fluorescence could be the result of structural re-arrangements in the FMN-binding region of mSOG during its encapsulation. Herein, aromatic amino acid residues (i.e. Trp) become exposed to the isoalloxazine ring of the FMN chromophore, leading to π–π stacking interactions that quench FMN fluorescence, an effect known to occur with other fluorescent flavoproteins. ^29^

### “On demand” generation of singlet oxygen from the light-activated Enc-mSOG nanoreactor

*Tm* encapsulin has multiple 3-4 Å sized surface pores that allow the flow of small metal ions (i.e. Fe^2+/3+^) into and out of its internal cavity. ^3, 13^ As depicted in Figure 2a, we hypothesized that molecular oxygen (O_2_) substrate can diffuse through the open surface pores of Enc-mSOG, enabling its interaction with the mSOG cargo within. Enc-mSOG can then be activated “on demand” with blue light to photoconvert O_2_ into ^1^O_2_. The highly reactive ^1^O_2_ product subsequently exits Enc-mSOG via its surface pores, allowing it to react with nearby molecules i.e. photosensitization.

**Fig. 2.**
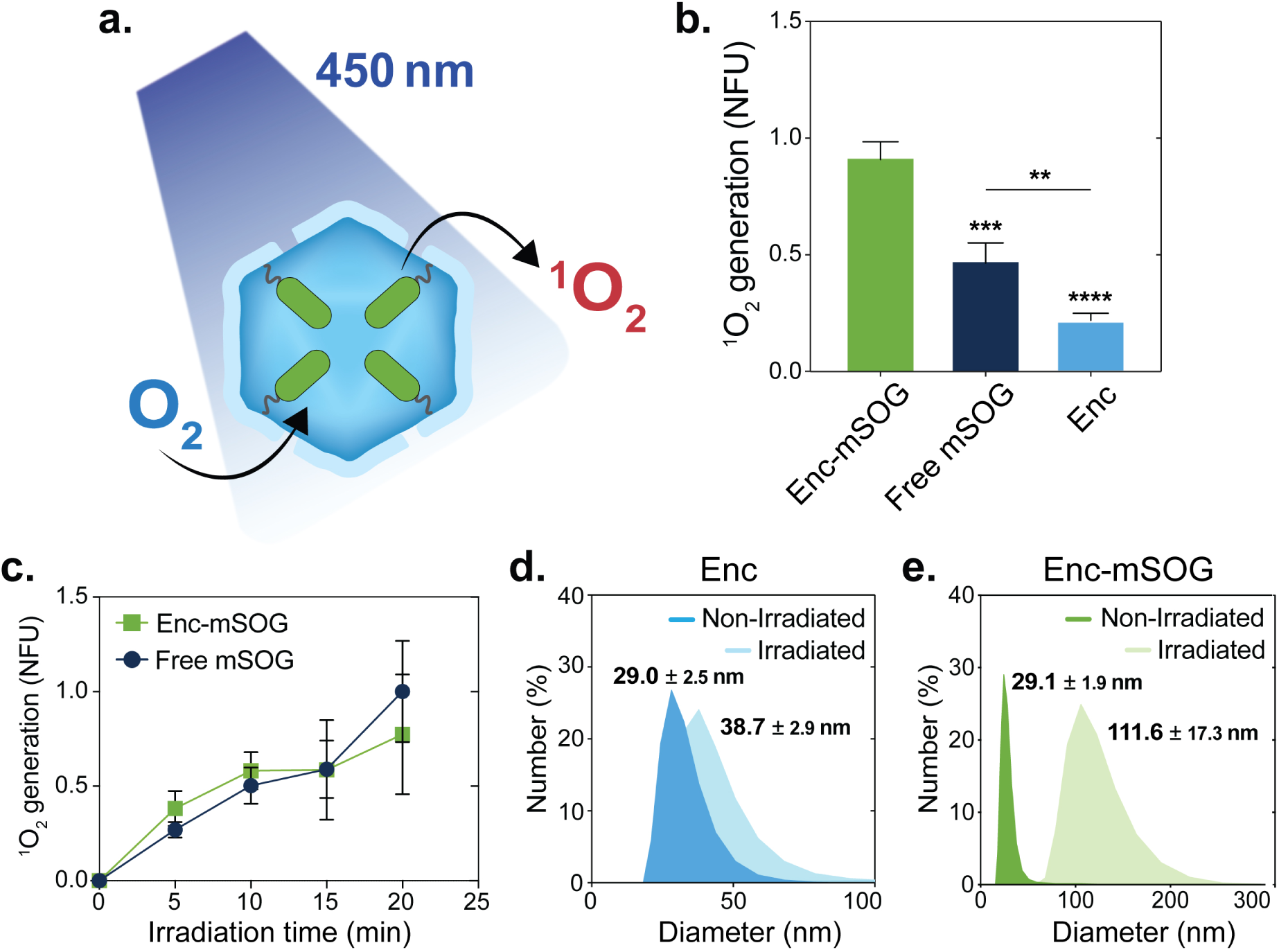
The “on demand” generation of singlet oxygen by light-activated Enc-mSOG. **(a)** Illustration showing the production of singlet oxygen (^1^O^2^) by a light-activated Enc-mSOG nanocompartment. In this process, molecular oxygen (O^2^) enters the internal cavity of the nanocompartment via open surface pores where it interacts with encapsulated mSOG cargo. Upon activation with blue laser light, the mSOG cargo converts O^2^ (substrate) into ^1^O_2_ (product), which subsequently diffuses out of the same open surface pores. **(b)** ^1^O_2_ generation from Enc-mSOG, free mSOG, and unloaded Enc upon irradiation with a blue laser (450 nm) at 55 mW/cm^2^ for 10 min. ^1^O_2_ production was determined by the fluorescence intensity of oxidized SOSG, (NFU). Error bars represent the mean ± standard deviation, one-way ANOVA, Tukey, n=3 (* p ≤ 0.05, ** p ≤0.01, *** p ≤ 0.001, **** p ≤ 0.0001). **(c)** ^1^O_2_ production by free and encapsulated mSOG after laser irradiation at 450 nm (55 mW/cm^2^) for 0, 5, 10, 15 and 20 min. ^1^O_2_ generation was measured using SOSG. It should be noted that the background ^1^O_2_ produced by unloaded Enc was subtracted from the total ^1^O_2_ produced by Enc-mSOG. Error bars represent the mean ± standard deviation n=3. **(d)** and **(e)** DLS-measured size distributions of unloaded Enc and Enc-mSOG before (Non-Irradiated) and after (Irradiated) laser excitation at 450 nm (55 mW/cm^2^) for 10 min.

To test this hypothesis, the capacity for light-activated Enc-mSOG to generate ^1^O_2_ in solution was measured using SOSG reagent. SOSG is selectively oxidized by ^1^O_2_ to emit green fluorescence (510-550 nm), the intensity of which is relative to the quantity of ^1^O_2_ produced. Enc-mSOG, free mSOG, and unloaded Enc were mixed with SOSG (in deuterated-HEPES buffer), and each sample irradiated with a blue laser for 10 min (450 nm). Afterwards, their fluorescence intensity was measured at 520 nm. Figure 2b shows that unloaded Enc is capable of producing low amounts of ^1^O_2_ upon laser excitation. Nevertheless, when compared to unloaded Enc, free and encapsulated mSOG generated 2.2-fold and 4.3-fold more ^1^O_2_, respectively. Unexpectedly, Enc-mSOG produced 1.9-fold more ^1^O_2_ generation than free mSOG, likely indicating that Enc has an additive effect when combined with mSOG’s own ^1^O_2_ generation. The relatively low quantities of ^1^O_2_ produced by unloaded Enc is likely due to the non-specific absorption of endogenous flavin molecules (e.g. FMN, flavin adenine dinucleotide (FAD), riboflavin) onto the Enc’s outer surface, which has been previously observed for *E. coli* produced *Tm* encapsulin. ^30^

In order to further characterize the functional effect encapsulation has on mSOG’s ^1^O2*-* generating capacity, the amounts of ^1^O_2_ produced by free and encapsulated mSOG were compared after different irradiation time periods (0, 5, 10, 15, 20 min) (Figure 2c). To account for background ^1^O_2_ from Enc, we subtracted this value from Enc-mSOG’s ^1^O_2_ generation at each time point. At all tested irradiation times, free and encapsulated mSOG produced similar amounts of ^1^O_2_ (Figure 2c), indicating that the encapsulation of mSOG and its subsequent loss of fluorescence intensity (Figure 1e) had no significant adverse effect on its ^1^O_2_-generating function. However, Enc-mSOG’s ^1^O_2_ production began to plateau at irradiation times longer than 10 min. Notably, without the subtraction of Enc’s background ^1^O_2_ production, Enc-mSOG was found to produce statistically greater amounts of ^1^O_2_ than free mSOG, thus confirming an additive effect when combining Enc and mSOG’s ^1^O_2_ generating capacities (Supplementary Figure 2).

The protein shells of encapsulins are considered robust nanostructures, exhibiting resilience against extreme pH, high temperatures and proteolytic degradation. ^1, 23, 25^ However, ^1^O_2_ is highly reactive against some amino acids (e.g. His, Tyr, Met, Cys) and can therefore cause problematic oxidative damage to protein structures. ^31^ To assess the physical effect of laser irradiation and ^1^O_2_ generation on the nanocompartments, we monitored changes to the structure and stability of unloaded Enc and Enc-mSOG after exposure to a blue laser (55 mW/cm^2^, 10 min). Following the irradiation of empty Enc, DLS measurements indicated a ∼31 % increase in its average hydrodynamic diameter from 29.0 to 38.7 nm, while TEM images revealed the presence of predominantly normal spherical nanocompartments with only a small proportion of large amorphous structures (Figure 2d and Supplementary Figure 3). Thus, laser irradiation alone had a minimal effect on the protein shell’s physical properties in accordance with a minimal ^1^O_2_ generation. In contrast, DLS measurements showed that irradiated Enc-mSOG enlarged ∼283.5 % from 29.1 to 111.6 nm (Figure 2e) and lost its monodispersity. Under TEM, a highly heterogeneous population was observed, consisting of enlarged nanocompartments and numerous bulky amorphous structures (Supplementary Figure 3). The loss of Enc-mSOG structural integrity and stability can be attributed to the light-induced activation of its mSOG cargo, which generates ^1^O_2_ that could severely damage its surrounding protein shell. ^*32*^ This is consistent with research by Zhen et al., in which ferritin protein nanocages were loaded with the potent chemical photosensitizer ZnF_16_Pc, and subsequently destroyed by ^1^O_2_ generated from the light-activated ZnF_16_Pc cargo. ^*33*^ Furthermore, the ^1^O_2_-mediated damage to Enc-mSOG macrostructure could explain why its photoconversion rate begins to plateau after exposure to more than 10 min laser irradiation (Figure 2c).

### Evaluating the photosensitizing function of Enc-mSOG in an in vitro model of photodynamic therapy (PDT)

*In vitro* studies have demonstrated miniSOG’s potential as a biological photosensitizer for PDT. ^34, 35^ To eliminate tumour cells, PDT relies on light-induced photosensitizers that convert intracellular oxygen into ROS, which damage cellular componentry and cause cell death. ^36^ As a proof-of-concept (Figure 3a), we decided to explore whether ROS produced by the light-activated Enc-mSOG nanoreactor was sufficient to trigger cellular photodynamic responses in an *in vitro* model of human lung cancer.

**Figure 3.**
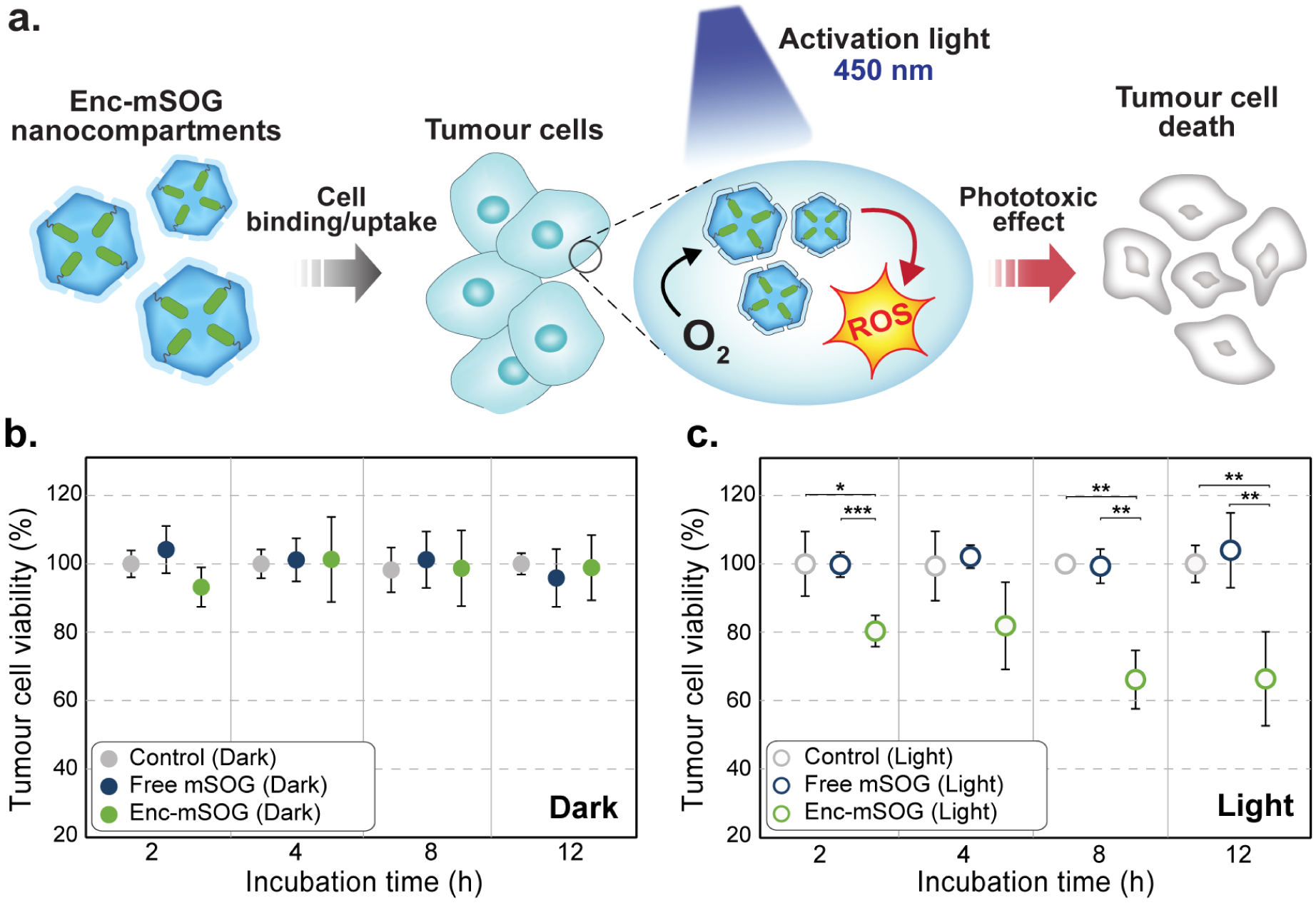
Testing the capacity of light-activated Enc-mSOG to induce photodynamic responses in an *in vitro* cancer model. **(a)** Schematic diagram showing the proposed delivery, activation and phototoxic effect of Enc-mSOG nanocompartments. Photosensitizing Enc-mSOG enters tumour cells via endocytosis. Upon photoexcitation with blue light, Enc-mSOG converts intracellular O_2_ into cytotoxic ROS (e.g. ^1^O_2_) that induces tumour cell death. **(b)** Cytotoxicity of free and encapsulated mSOG: viability of A549 cells after incubation without (control) or with mSOG or Enc-mSOG for different times (2, 4, 8, and 12 h) in the dark. Cell viability was subsequently determined by MTT assay. Error bars represent the mean ± standard deviation (p≤ 0.05), one-way ANOVA, Dunnett T3, n=6 from two independent experiments. **(c)** Phototoxicity of free and encapsulated mSOG: Viability of A549 cells incubated without (control) or with free mSOG or Enc-mSOG for different times (2, 4, 8, and 12 h) in the dark, followed by blue laser irradiation at 450 nm (55 mW/cm^2^) for 10 min. Cell viability was quantified via MTT assay. Error bars represent the mean ± standard deviation (*p≤ 0.05, **p≤0.01), one-way ANOVA, Dunnett T3, n=6 from two independent experiments.

First, the cytotoxicity of unloaded Enc, free mSOG and Enc-mSOG was assessed in dark conditions. Previous work by Deyev demonstrated that miniSOG tagged with a cancer-specific antibody mediated targeted PDT *in vitro*, exerting its maximal phototoxic effect against cancer cells at a concentration of 500 nM. ^34^ Accordingly, A549 human lung cancer cells were pre-incubated with 4.8 mM of unloaded Enc and 500 nM mSOG or Enc-mSOG for increasing periods of time (2, 4, 8 and 12 h) in the dark, after which viability was measured by MTT assay. In the absence of light activation, free mSOG and Enc-mSOG showed no significant cytotoxicity when compared to untreated cells (Figure 3b). This was also the found to be the case for unloaded Enc (Supplementary Figure 5a).

Next, we sought to evaluate the cytotoxicity of unloaded Enc and free and encapsulated mSOG in conjunction with blue-light irradiation i.e. phototoxicity. Initially, the effect of laser irradiation on live cells was assessed by exposing A549 cells to a blue laser (55 mW/cm^2^). An irradiation time of up to 10 min had no significant effect on cell viability (Supplementary Figure 4) and was therefore used to study *in vitro* phototoxicity. Herein, A549 cells were treated using the same conditions outlined above, followed by 10 min blue-light irradiation, with cell viability then determined via MTT assay. As shown in Figure 3c, A549 cells showed no significant changes in viability after pre-incubation with free mSOG (2, 4, 8 and 12 h) and blue-light irradiation. This is consistent with reports in which unmodified free miniSOG was unable to efficiently bind or enter cancer cells and exert its phototoxic effect ^34^. Likewise, unloaded Enc elicited no significant decrease in cell viability upon light irradiation (Supplementary Figure 5b), suggesting that ^1^O_2_ generated by Enc alone is too low to be cytotoxic. In contrast, cells pre-incubated with Enc-mSOG (2, 4, 8 and 12 h) and exposed to laser irradiation exhibited a decrease in viability for each treatment tested. For instance, the viability of cells pre-incubated with Enc-mSOG for 8 or 12 h were significantly reduced by ∼34%. Thus, Enc serves as a viable nanocarrier for mSOG, localizing it in tumour cell membranes and interiors, where it can be triggered by light to induce oxidative damage that lowers cell viability.

## Conclusion

In summary, we reversed the oxidative stress response functions of *Tm* encapsulin, programming it to instead serve as a light-responsive encapsulin nanoreactor that generates ROS “on demand” and triggers photosensitization reactions. To realise this objective, the protein photosensitizer miniSOG was tagged with the *Tm* ESig and selectively packaged into *Tm* encapsulin, resulting in a self-assembled Enc-mSOG nanoreactor that showed similar structural properties to unloaded nanocompartments.

Upon encapsulation, mSOG retained its fluorescence spectra but lost its fluorescence intensity, potentially due to structural rearrangements. Importantly, we found that confinement had no effect on mSOG’s ability to photoconvert O_2_ to ^1^O_2_ under blue-light excitation. Specifically, free and encapsulated mSOG generated comparable amounts of ^1^O_2_, and empty Enc produced small quantities of ^1^O_2_. Enc-mSOG’s total ^1^O_2_ production confirmed an additive effect between mSOG cargo and Enc, a positive attribute that provides a unique advantage over the use of free mSOG alone. Additionally, these findings infer the adsorption of endogenous photosensitizing flavins (e.g. FMN and FAD) onto *Tm* encapsulin, offering insight into its physiological function.

After prolonged blue-light exposure, Enc-mSOG began to show signs of structural deterioration, which coincided with a slow plateau in its photoconversion rate. These adverse effects were likely caused by the rapid generation of destructive ^1^O_2_ inside Enc-mSOG’s protein shell. ^31^ Given their structural adaptability, encapsulin protein shells could be re-engineered to become more resilient towards ROS. For instance, residues prone to ^1^O_2_ oxidation could be substituted with ^1^O_2_-insensitive residues, while surface pores could be enlarged to avoid product accumulation and increase substrate turnover. ^37, 38^

As a proof of concept, light-activated Enc-mSOG triggered photosensitized oxidation reactions that significantly lowered the viability of lung cancer cells. To our knowledge, this is the first time encapsulins have been used to deliver protein cargo in a therapeutic manner. Nevertheless, higher intracellular doses of Enc-mSOG will be required to maximize its efficacy in future PDT studies. This could be accomplished by engineering Enc-mSOG’s external surface to display targeting ligands (e.g. peptides and antibodies) that increase cancer cell uptake. In an example of this approach, *Tm* encapsulin was modified to co-display hepatocellular carcinoma (HCC)-targeting peptides and anticancer drugs, resulting in the effective delivery of drugs into HCC cells. ^23^

Due to the high spatial and temporal resolution offered by light, we expect that the novel Enc-mSOG nanoreactor can be selectively activated to initiate and/or modulate other ROS-sensitive processes with technological, biological and therapeutic relevance. Ultimately, this work illustrates the remarkable versatility of encapsulin nanoreactors, paving the way towards the establishment of new techniques and mechanisms that can more precisely control their activities.

## Acknowledgments

D.D. is supported by an international Macquarie University Research Excellence Scholarship (iMQRES), Sydney Vital Research Scholar Award, and the Commonwealth Scientific and Industrial Research Organisation (CSIRO) PhD Scholarship Program in Synthetic Biology. A.C. is supported by a Cancer Institute New South Wales Early Career Fellowship (Project Number: ECF171114), the Cancer Australia Priority-driven Collaborative Cancer Research Scheme (Project Number: 1182082), and the Australian Research Council (CE140100003).

## Author contributions

D.D co-designed the research, generated all nanocompartment constructs, conducted all physical and functional characterization work, performed data analysis and wrote the manuscript. X.V. constructed the laser set-up, supported laser irradiation experiments and revised the manuscript. A.S. supervised the project and revised the manuscript. A.C. conceptualized and co-designed the study, supervised the project and wrote the manuscript.

