## Supplementary Information for "Bioengineering a light-responsive encapsulin nanoreactor: a potential tool for photodynamic therapy"

**Supplementary Table 1.** Expression plasmids constructed for this study.

| Plasmids | Description | Protein ID |
|---|---|---|
| pACYC-Duet-1_Enc | Encapsulin from *Thermotoga maritima (Tm)*<br>138-GGGGGGHHHHHHGGGGGG-139 | UniProt: Q9WZP3 |
| - | *Tm* Encapsulation signal peptide (ESig) | UniProt: R4NZH3 |
| - | miniSOG: fluorescent flavoprotein protein engineered from *Arabidopsis thaliana* phototropin-2. | Phototropin-2<br>UniProt: P93025 |
| pPETDuet-1_mSOG-ESig | miniSOG-GGSENTGGDLGIRKL | |

**Supplementary Table 2.** Optimized conditions for the recombinant production of all proteins in *E. coli.*

| Protein(s) | Plasmid(s) | Antibiotic(s) | IPTG (mM) | Induction temperature (°C) | Induction time (h) |
|---|---|---|---|---|---|
| Enc | pACYC-Duet-1_Enc | Chloramphenicol | 0.1 | 37 | 4-6 |
| mSOG | pPETDuet-1_mSOG-ESig | Carbenicillin | | | |
| Enc-mSOG | pACYC-Duet-1_Enc<br>pPETDuet-1_mSOG-ESig | Chloramphenicol<br>Carbenicillin | 0.05 | 30 | 18-20 |

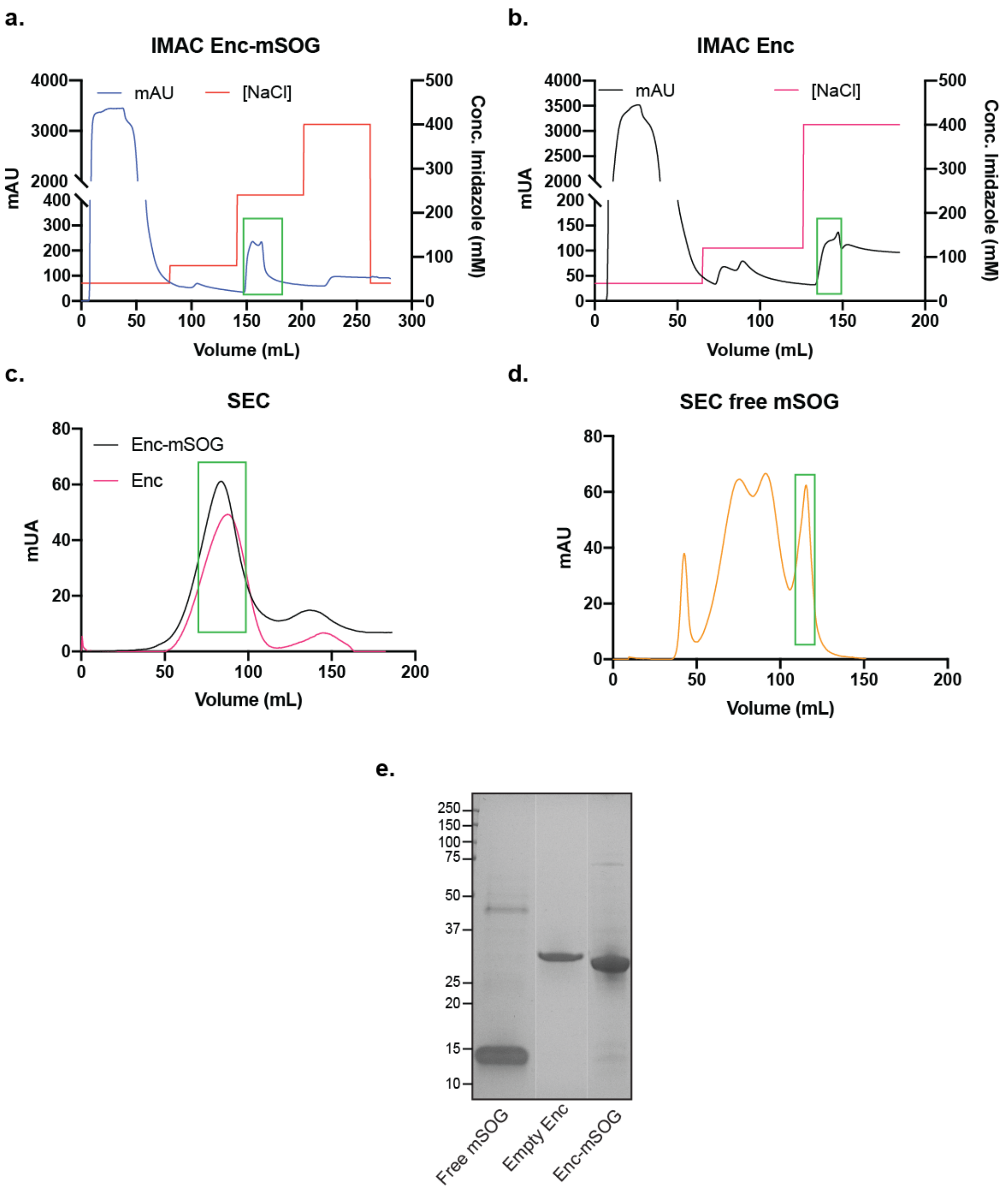


**Supplementary Figure 1. Example chromatograms of recombinant protein purifications.** Nickel-immobilized metal-affinity chromatography (IMAC) of **(a)** Enc-mSOG and **(b)** empty Enc. The green square highlights the peak corresponding to the protein of interest. **c.** Size exclusion chromatography (SEC) of IMAC-purified Enc-mSOG (pink line) or empty Enc (black line). The protein of interest (green square) elutes between 70-95 mL. **d.** SEC of partially purified ESig-tagged mSOG (free mSOG). The protein of interest (green square) elutes between 106-120 mL. **e.** Coomassie-stained SDS-PAGE showing the purification and co-purification of the $Enc_{Subunit}$ (~31.9 KDa) and mSOG (~14.4 KDa) cargo. The proteins showed a purity of 82% for free mSOG, 99% for Enc and 93% for Enc-mSOG.

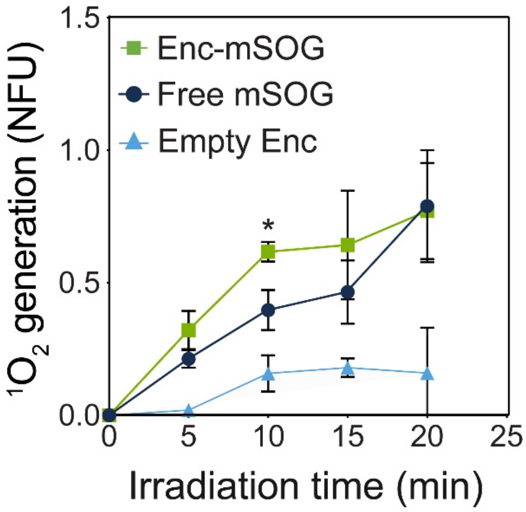


**Supplementary Figure 2. Total $^1O_2$ production from unloaded Enc, and free or encapsulated mSOG.** All protein constructions were laser irradiated at 450 nm (55 mW/cm$^2$) for 0, 5, 10, 15 or 20 min. $^1O_2$ generated was measured using SOSG and given as normalized fluorescence units (NFU). Error bars represent the mean ± standard deviation two-way ANOVA, Holm-Sidak, n=3 (* $p \leq 0.05$). At 10 min of irradiation Enc-mSOG's $^1O_2$ generation was significantly greater than free mSOG, indicating an additive effect between the Enc and mSOG's $^1O_2$ generating capacities.

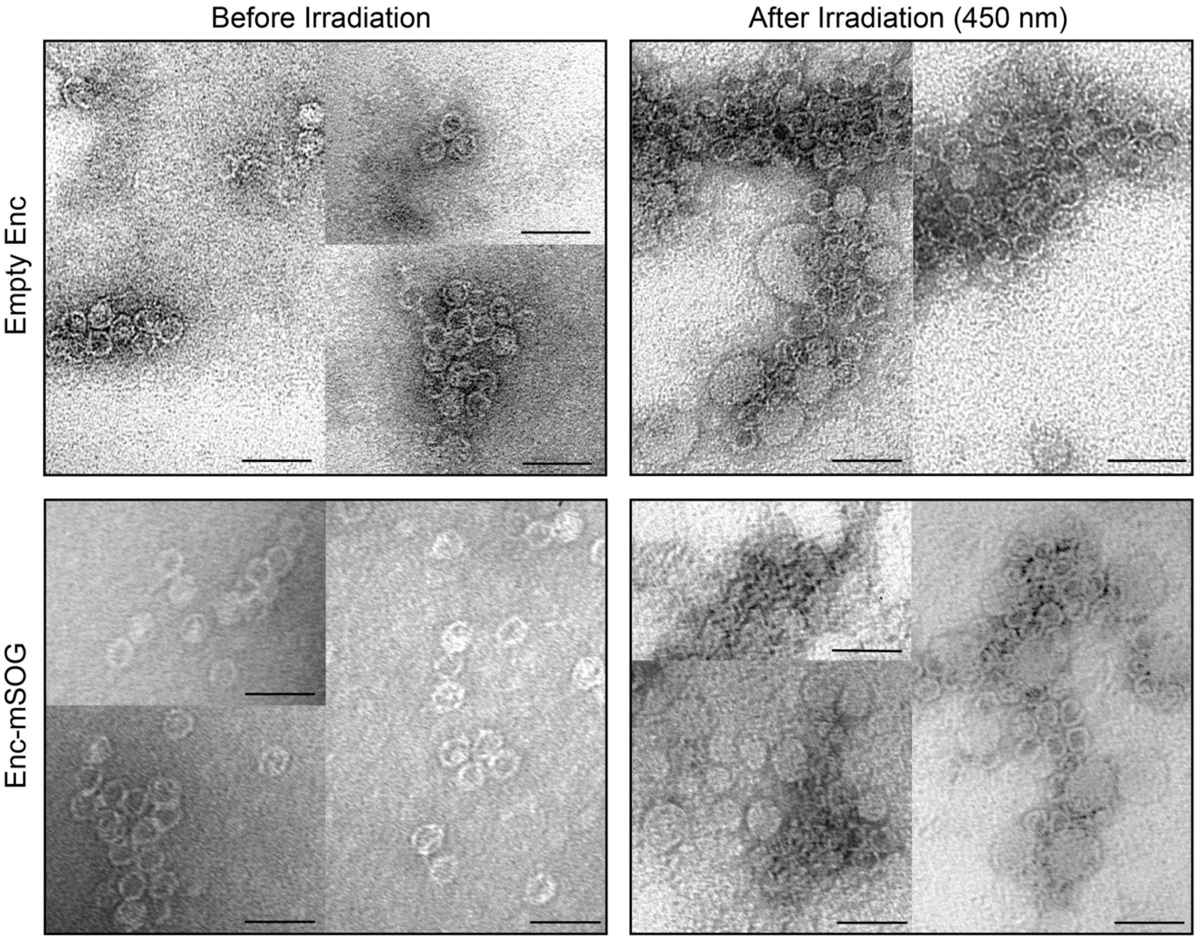


**Supplementary Figure 3. Effect of laser irradiation and $^1O_2$ generation on encapsulin structure**. TEM images showing **(Top panel)** empty Enc and **(Bottom panel)** Enc-mSOG before and after laser irradiation at 450 nm (55 mW/cm2) for 10 min (Scale bars = 50 nm).

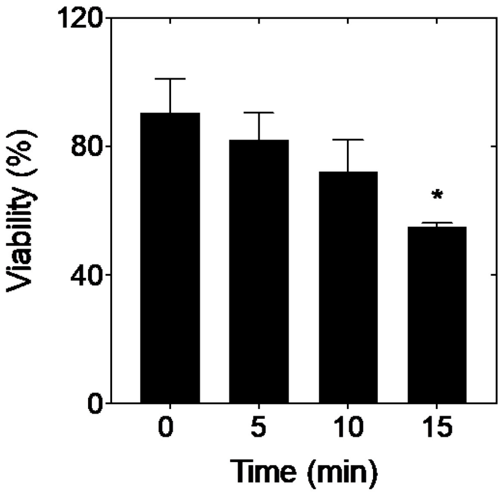


**Supplementary Figure 4. Effect of laser irradiation on A549 lung cancer cells.** Cell viability of A549 cells after laser irradiation at 450 nm (55 mW/cm2) for different time periods (0-15 min). Error bars represent the mean ± standard deviation (p≤ 0.05), one-way ANOVA, Tukey, n≥3.

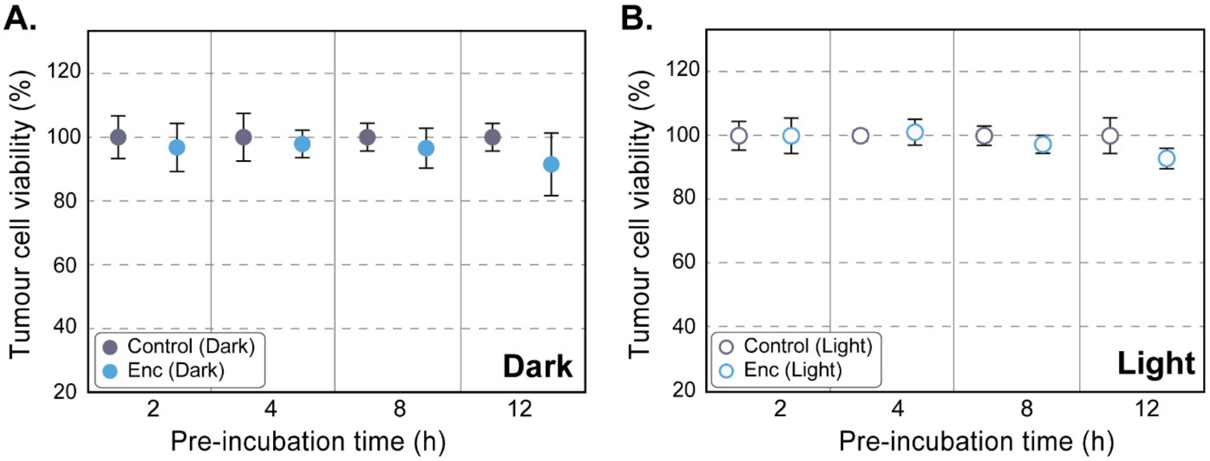


**Supplementary Figure 5. Evaluating the effect of unloaded Enc on A549 cancer cells. A.** Cytotoxicity of unloaded Enc: viability of A549 cells after incubation without (control) or with Enc for different times (2, 4, 8, and 12 h) in the dark. Cell viability was subsequently determined by MTT assay. Error bars represent the mean ± standard deviation (p≤ 0.05), one-way ANOVA, Dunnett T3, n=6 from two independent experiments. **B.** Phototoxicity of unloaded Enc: Viability of A549 cells incubated without (control) or with Enc for different times (2, 4, 8, and 12 h) in the dark, followed by blue laser irradiation at 450 nm (55 mW/cm$^2$) for 10 min. Cell viability was quantified via MTT assay. Error bars represent the mean ± standard deviation (*p≤ 0.05, **p≤0.01), one-way ANOVA, Dunnett T3, n=6 from two independent experiments.
